# Identification of single nucleotide variants using position-specific error estimation in deep sequencing data

**DOI:** 10.1101/475947

**Authors:** Dimitrios Kleftogiannis, Marco Punta, Anuradha Jayaram, Shahneen Sandhu, Stephen Q. Wong, Delila Gasi Tandefelt, Vincenza Conteduca, Daniel Wetterskog, Gerhardt Attard, Stefano Lise

**Affiliations:** Centre for Evolution and Cancer, The Institute of Cancer Research, London, UK; UCL Cancer Institute, University College London, London, UK; Peter MacCallum Cancer Centre and University of Melbourne, Melbourne, Victoria, Australia; Department of Urology, Sahlgrenska Academy, University of Gothenburg, Gothenburg, Sweden; Department of Medical Oncology, Istituto Scientifico Romagnolo per lo Studio e la Cura dei Tumori (IRST) IRCCS, Meldola, 47014, Italy; Genome Institute of Singapore (GIS), Agency for Science Technology and Research (ASTAR), Singapore, 138672, Singapore

**Keywords:** Next generation sequencing (NGS), cancer genomics, variant calling, deep sequencing, targeted sequencing, Ion Torrent, liquid biopsies, error correction

## Abstract

**Background:** Targeted deep sequencing is a highly effective technology to identify known and novel single nucleotide variants (SNVs) with many applications in translational medicine, disease monitoring and cancer profiling. However, identification of SNVs using deep sequencing data is a challenging computational problem as different sequencing artifacts limit the analytical sensitivity of SNV detection, especially at low variant allele frequencies (VAFs).

**Methods:** To address the problem of relatively high noise levels in amplicon-based deep sequencing data (e.g. with the Ion AmpliSeq technology) in the context of SNV calling, we have developed a new bioinformatics tool called AmpliSolve. AmpliSolve uses a set of normal samples to model position-specific, strand-specific and nucleotide-specific background artifacts (noise), and deploys a Poisson model-based statistical framework for SNV detection.

**Results:** Our tests on both synthetic and real data indicate that AmpliSolve achieves a good trade-off between precision and sensitivity, even at VAF below 5% and as low as 1%. We further validate AmpliSolve by applying it to the detection of SNVs in 96 circulating tumor DNA samples at three clinically relevant genomic positions and compare the results to digital droplet PCR experiments.

**Conclusions:** AmpliSolve is a new tool for *in-silico* estimation of background noise and for detection of low frequency SNVs in targeted deep sequencing data. Although AmpliSolve has been specifically designed for and tested on amplicon-based libraries sequenced with the Ion Torrent platform it can, in principle, be applied to other sequencing platforms as well. AmpliSolve is freely available at https://github.com/dkleftogi/AmpliSolve.

## Background

Targeted next-generation sequencing (NGS) is a powerful technology to identify known and novel variants in selected genomic regions of interest [1]. It allows achieving high coverage levels (i.e., higher than 1000x) and, in principle, to confidently identify variants even when they occur at low allele frequencies. This is particularly important in cancer research and has many clinical applications, e.g. in relation to disease management. Typically, tumors are heterogeneous consisting of multiple clones and sub-clones the relative abundance of which can change over time depending on several factors, including treatment [2]. Identification of low frequency mutations is clinically relevant, among other reasons, for early diagnosis, disease monitoring and timely detection of the emergence of resistance clones under treatment [3].

Over the past years, it has been established that cancer patients’ circulating free DNA (cfDNA) contains tumor-derived DNA fragments (ctDNA) that can be used as an alternative to solid biopsies in clinical settings [4]. However, identifying cancer-specific mutations in liquid biopsy samples is challenging, as the relative proportion of ctDNA in cfDNA can be low, especially at cancer’s early stages. There are also several sources of sequencing errors including PCR artifacts, often reaching up to 1% Variant Allele Frequency (VAF), that reduce further the analytical sensitivity for detecting cancer-associated mutations [4]. Error correction techniques can be incorporated into NGS assays enabling ultra-sensitive single nucleotide variant (SNV) detection (VAF ~ 0.1%) but at a significant extra cost [5,6]. Thus, there is a need to reliably detect SNVs in more conventional deep sequencing data.

*In silico* identification of SNVs from NGS data is a well-studied problem [7,8]. However, the majority of existing variant calling programs have been designed for whole-exome and whole-genome experiments sequenced at coverage of approximately 30x to 100x. At the same time, available variant calling software for targeted deep sequencing experiments have been typically developed for and tested on Illumina data [9].

Compared to Illumina, Ion Torrent sequencing has a higher per base error rate and an associated lower accuracy in identifying mutations [10, 11]. However, it has the advantage of requiring lower amounts of input DNA and it offers both reduced cost and turnaround time. Thus, it is a cost-effective strategy for screening large cohorts of patients and it is particularly suited for point-of-care clinical applications [1], for example in conjunction with the Ion AmpliSeq Cancer Hotspot Panel. Given its translational potential, there is a real need to improve the variant calling workflow and recently a number of methods have been developed to deal specifically with Ion Torrent data [12, 13, 14].

Here we introduce AmpliSolve, a new bioinformatics method to detect SNVs in targeted deep sequencing data. It combines *in-silico* background error estimation with statistical modeling and it is particularly suited to deal with data of comparatively high noise levels, similar to the ones produced by the Ion AmpliSeq library preparation. In order to estimate background noise levels per position, strand and nucleotide substitution, AmpliSolve takes as input deep-sequencing data from a set of normal samples. This information is then fed to a Poisson model for the identification of SNVs. Experimental results using normal samples (self-consistency test), synthetic variants and clinical data sequenced with a custom Ion AmpliSeq gene panel, demonstrate that AmpliSolve achieves a good trade-off between precision and sensitivity, even for VAF values below 5% and as low as 1%.

## Methods

### Method overview

AmpliSolve consists of two main programs written in C++: AmpliSolveErrorEstimation and AmpliSolveVariantCalling. AmpliSolveErrorEstimation requires the availability of a set of normal samples processed with the deep sequencing platform and panel of choice. Here, we focus on the Ion Torrent Personal Genome Machine (PGM) and a custom AmpliSeq panel, a technology known to have relatively high rates of sequencing error compared to others. The program uses the normal samples to infer position-specific, nucleotide specific and strand-specific background sequencing error levels (noise) across the targeted regions. Execution of AmpliSolveErrorEstimation is performed only once per panel design. Error estimates are then used as input to the AmpliSolveVariantCalling program for SNVs’ detection. The procedures for *in-silico* noise estimation and SNV identification are described below. In Figure 1 we present a graphical overview of the AmpliSolve computational workflow.

**Figure 1.**
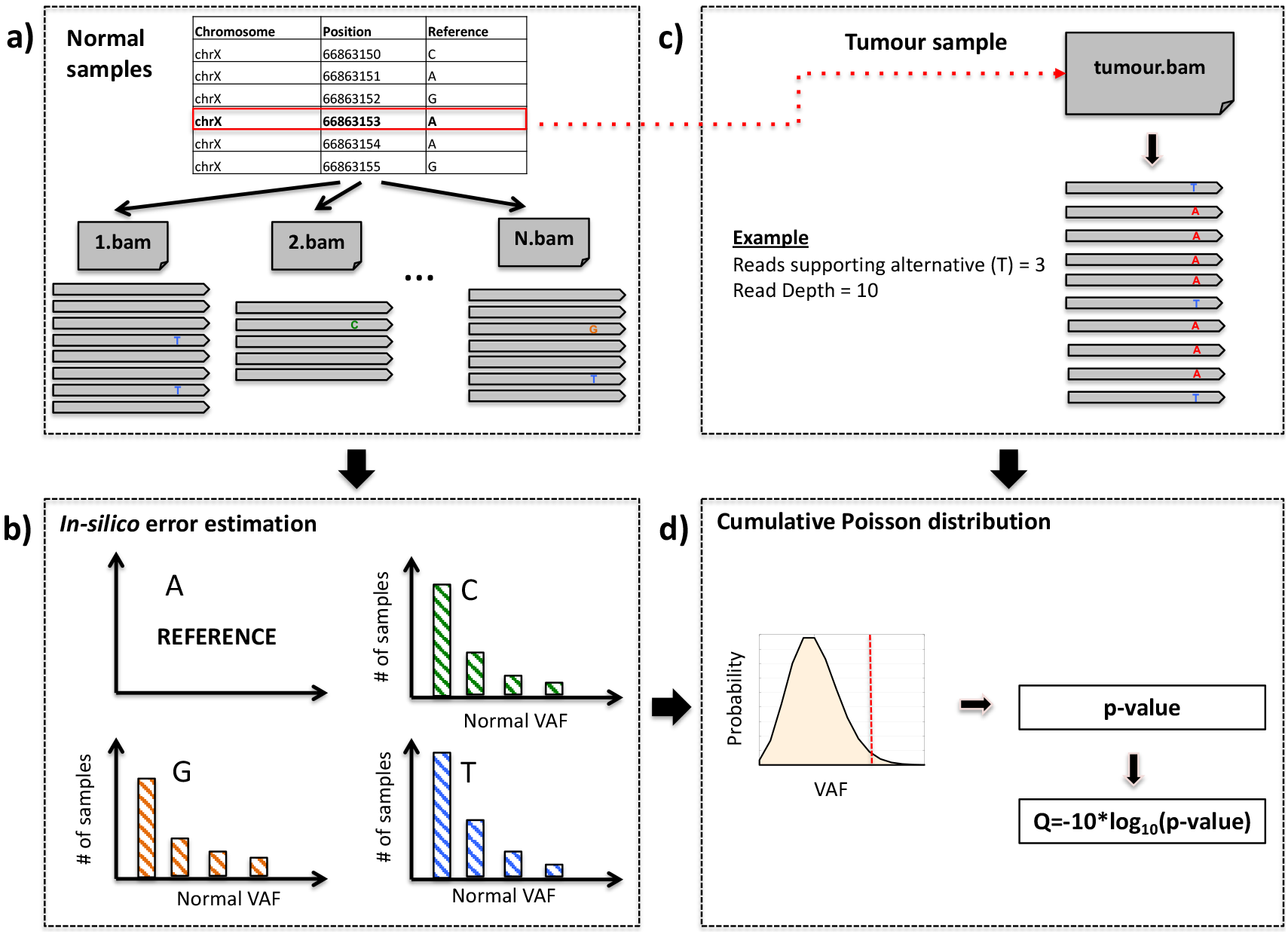
Graphical representation of AmpliSolve’s workflow for estimating the noise levels and detecting SNVs. The workflow comprises the following steps: a) Screening the available normal samples to identify reads supporting alleles other than the reference. b) Error estimation per position, per nucleotide and per strand for all positions in the gene panel based on the distribution of alternative allele counts in (a); only alternative counts corresponding to VAF<5% are taken into consideration; c) For each genomic position in a tumor sample, the method identifies the total coverage of the position and the number of reads supporting the alternative alleles, if any. d) Given the information from steps b) and c) the method applies a Poisson distribution-based model to compute the p-value that the variant (red line) is real. This p-value is then transformed to a quality score that is used by AmpliSolve together with additional quality criteria to identify SNVs.

### *In-silico* identification of the background sequencing error

Our strategy for estimating background error levels, implemented in the AmpliSolveErrorEstimation program, is based on the assumption that alternative alleles observed at VAF<5% in normal samples are, in the majority of cases, the result of sequencing errors (see Figure S1 for the distribution of non-reference allele frequencies in normal samples showing the separation between heterozygous germline variants and lower frequency ‘noise’ variants). Accordingly, we utilize a set of normal samples to estimate background noise in our custom panel. Notably, we estimate error levels separately for each genomic position, each nucleotide (alternative allele) and each of the two (forward and reverse) strands. In particular, for each genomic position we generate six error estimates (i.e. two each for the three alternative alleles). Error estimates are fed to a Poisson model, which is then used to calculate the p-value of the observed substitutions representing true variants versus them being noise. The detailed implementation is as follows. We first extract “raw” counts for each position, alternative allele and strand from the BAM files [15] of a set of N normal samples using the ASEQ software [16]. We run ASEQ with the quality parameters suggested by the authors of a previous study based on Ion AmpliSeq data [17], namely: minimum base quality = 20, minimum read quality = 20 and minimum read coverage = 20. At every genomic position, we estimate the background error *s* separately for each alternative allele *α* and strand (+ or −) by calculating the fraction of reads carrying the alternative allele on a given strand across all normal samples. More specifically we use the following formula:

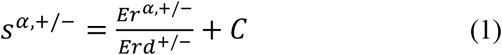

with

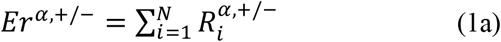

and

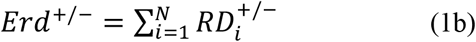

We denote with R_i_^α,+^ and R_i_^α,−^ the number of reads supporting the alternative allele α on the forward and reverse strand, respectively, in normal sample *i*. We denote with RD_i_^+^ and RD_i_^−^ the total number of reads (read depth) at the genomic position of interest on the forward and reverse strand, respectively, in normal sample *i*. Summations are taken over all normal samples utilized for the error estimation. C in equation (1) is a constant pseudo-count parameter that is introduced to mitigate the problem of positions in which the alternative allele read count might be underestimated (e.g. due to a relatively low read depth at a given position in the normal samples). In the Results section we test values of C in the range from 10^−5^ to 2·10^−2^.

When calculating the summations in (1a) and (1b), we apply two filters that aim to increase the quality of the error estimation at specific positions and for specific alternative alleles *α* at that position. First, at a given position, samples for which an alternative allele *α* has VAF > 5% are not considered at that position. This is because a frequency greater than 5% is likely to indicate, in that sample, either the presence of a real variant (i.e. a single nucleotide polymorphism) or a particularly noisy ‘read-out’. Second, samples that at a given position have coverage lower than a predefined threshold either on the forward or on the reverse strand are not considered for computing *Er*^*α*,+/−^ (1a) and *Erd*^+/−^ (1b) for any allele *α* at that position. In the following we use a threshold of 100 reads which, in our case, typically excludes 5% of sites per sample (see Figure S2); however, this parameter can be adjusted depending on the study design. After applying these filters, positions and alternative alleles for which 2/3 or more of the normal samples cannot be used for calculating the summations in (1a) and (1b) are considered non-callable. We note that among non-callable cases there may be positions with alleles that are either frequent in the population or simply over-represented in the specific set of normal samples used for the error estimation. However, given that AmpliSolve main goal is the identification of somatic mutations this does not constitute a major limitation.

### SNV detection using a cumulative Poisson distribution

Given a sample of interest, for every alternative allele *α* featuring a non-zero strand-specific (+ or −) variant read count *k*^*α*,+/−^, the AmpliSolveVariantCalling program uses a Poisson model to calculate the probability that *k*^*α*,+/−^ or more variant reads are produced by sequencing errors, i.e. the p-value. Only positions that, in the sample of interest, have read depth on each strand higher than a pre-assigned threshold *RD*_*min*_ are considered (in the following, we set *RD*_*min*_ = 100 unless otherwise specified). At these positions, the calculated p-value is a function of the normal sample-based sequencing error *s*^*α*,+/−^ from the previous section and of both the number of variant reads *k*^*α*,+/−^ and the strand-specific read depth *K*^+/−^ in the sample of interest (*K*^+/−^ > *RD*_*min*_). In particular:

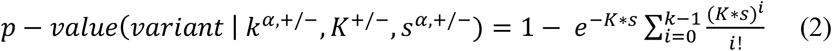

Where, for better readability, on the right side of the equation we have omitted all *α* symbols for k and s, as well as, +/− symbols for k, K and s. We observe that *K*\**s* is the expected number of random substitutions for a depth of coverage *K* or the mean of the Poisson distribution. Note that p-values are not corrected for multiple testing. The strand-specific p-values are finally converted to quality scores using the formula *Q*=−*10*\**log10(p-value)*. In its output, for all positions in the panel carrying substitutions with Q score equal to or greater than 5 on both strands, AmpliSolve reports the average Q score between the two strands. *Vice versa*, positions with no substitutions or with substitutions with associated Q score lower than 5 on one or both strands are not reported in AmpliSolve’s output. All reported SNVs are further tested for and potentially assigned one or more of the following warning flags:

a. ‘LowQ’ if the Q score is lower than 20 in at least one of the two strands.
b. ‘LowSupportingReads’ if the SNV is supported by less than 5 reads per strand in the tumor samples being analysed.
c. ‘AmpliconEdge’ if the SNV is located within overlapping amplicon edge regions, which may result in sequencing artifacts.
d. ‘StrandBias’ if the SNV is associated to a strand-bias. We apply Fisher’s exact test to each SNV under the null hypothesis that the number of forward and reverse reads supporting the variant should be proportional to the total number of reads sequenced in the forward and reverse strands, respectively. The flag is assigned to substitutions for which the p-value of the Fisher’s exact test is lower than a pre-defined value *SB*_*th*_. In the following we set *SB*_*th*_ = 0.05 (unless otherwise specified).
e. ‘HomoPolymerRegion’ if the SNV is located within a homopolymer region using the same criteria as in [18].
f. ‘PositionWithHighNoise’ if the SNV is supported by more than 5 reads per strand but the associated VAF is lower than the maximum VAF at this position across all normal samples in the training set.

If no warning is issued, AmpliSolve assigns a ‘PASS’ flag to the SNV.

### Performance measures

To assess AmpliSolve’s success in detecting SNVs, we use a number of performance metrics:

1. Sensitivity or True Positive Rate (TPR) = TP / (TP+FN)
2. Precision or Positive Predictive Value (PPV)= TP/(TP+FP)
3. False Discovery Rate (FDR) = 1-PPV=FP / (FP+TP)
4. Harmonic mean of Precision and Sensitivity (F1) = 2*TP / (2*TP +FP + FN)

Where, TP is the number of True Positive predictions, FN is the number of False Negative predictions and FP is the number of False Positive predictions.

### Clinical data used in this study

For the development and evaluation of AmpliSolve, we have access to an extensive collection of clinical samples from castration-resistant prostate cancer (CRPC) patients, part of which had been already presented in previous publications [17,19,20,21]. The collection comprises 184 germline samples (white blood cells, buccal swabs or saliva) and more than 450 liquid biopsy plasma samples (note that for some patients there are multiple liquid biopsies and a small minority of liquid biopsy samples has no matched normal). In practice for this study, we rely on all 184 normal samples but only use 96 liquid biopsy samples for which results from digital droplet PCR (ddPCR) assays are available (see below). For 5 additional patients we have access to 10 solid tumor samples from metastatic sites (1, 2 and 3 samples from respectively 1, 3 and 1 patients) and their associated 5 germline samples. For the available samples, we have the following data:

a. For all samples (germlines, liquid biopsies and solid tumors), we have Ion Torrent sequencing data obtained using a custom Ion AmpliSeq panel of 367 amplicons spanning 40,814 genomic positions at around 1000-1500x coverage. The panel targets both intronic and exonic regions in chromosomes 8, 10, 14, 17, 21 and X including commonly aberrated genes such as *PTEN*, *CYP17A1*, *FOXA1*, *TP53*, *SPOP* as well as the androgen receptor (*AR*) gene, which is one of the main drivers of CRPC, and the drug target *CYP17A1*. More details about the sequencing protocol, data processing and additional information about the application of our custom Ion AmpliSeq panel in CRPC diagnostic studies can be found in [17] and [19]. These papers also include a description of a variant caller that we used as starting point for developing AmpliSolve. We call variants in these Ion AmpliSeq data with our program AmpliSolve.
b. For the 10 solid tumor samples and 5 matched germline samples, in addition to Ion Torrent data, we have Illumina Whole Genome Sequencing (WGS) data at around 80-100x (tumor) and 30x (germline) coverage. We call variants in WGS data according to a previously established pipeline [22], which we describe in the next section.
c. For 96 liquid biopsy samples, we have results from ddPCR assays to screen 3 clinically relevant SNVs in the *AR* gene. These SNVs have been linked to resistance to targeted therapy in CRPC patients, namely: 2105T>A (p.L702H), 2226G->T (p.W742C) and 2632A>G (p.T878A). ddPCR in the plasma samples was performed using 2-4 ng of DNA, using Life Technologies Custom Taqman snp genotyping assay (product codes AH0JFRC, C_175239649_10 and C_175239651_10, respectively). Following droplet generation (AutoDg, Bio-Rad) and PCR, samples were run on the Bio-Rad QX200 droplet reader and analyzed using the QuantaSoft software.

### WGS variant calling pipeline

We used Illumina WGS data to generate a benchmark set of calls (“ground-truth”) against which AmpliSolve performance is evaluated. WGS data have been processed using standard tools, such as Skewer [23] for adapter trimming, BWA-MEM [24] for mapping and Picard [25] for duplicate removal. In order to call SNVs we run a previously developed pipeline [22] that utilizes jointly Mutect [26] and Platypus [18] (throughout the manuscript this pipeline is denoted as *MutPlat*). Briefly, we first run Mutect (default parameters) on each paired tumor-normal samples. Then, we use Mutect’s calls as priors for Platypus and jointly call variants on all tumors and matched normal samples of a patient (further details are provided in Additional file 1: Supplementary Methods). Our ground-truth set of calls consists of both germline and somatic mutations extracted as explained below.

Germline variants are identified as those variants called in the normal (GT=0/1 or 1/1) and that, additionally, have either a PASS filter flag or don’t have a PASS filter flag (could have e.g. ‘badReads’) but are present in 1000 genomes (phase 3 release) [27]. For AmpliSolve validation purposes we consider only tumor samples but include both germline and somatic SNVs. By including germline SNVs, in particular, we are able to test a higher number of low VAF mutations than would be possible when considering only somatic mutations. This is due to a combination of somatic deletions and germline DNA contamination in the tumor samples. Indeed, while somatic deletions cause loss of some germline SNPs in tumor DNA, germline DNA contamination (i.e. <100% tumor purity) means that these mutations may still be present in the tumor samples, albeit with a lower VAF. Note that if somatic deletions occur in high tumor purity samples and/or the sequencing coverage is not deep enough, germline variants may have no supporting read at all in the WGS tumor data. We keep also these limiting cases as part of our ground-truth set of variants as they might be (and sometimes are) detectable in the targeted Ion AmpliSeq data.

To call a somatic SNV we require all of the following criteria to be met: i) Platypus filter: PASS, alleleBias, Q20, QD, SC or HapScore, ii) at least 3 reads supporting the variant in the tumor, iii) at least 10 reads covering the position in the germline and no support for the variant in the germline (NV=0 and genotype GT= 0/0), iv) SNV not present in the 1000 genomes database.

### SNV callers tested for comparison

On WGS and ctDNA sample, we compare AmpliSolve to SiNVICT [12], a tool that has been shown to be effective in detecting mutations at very low VAF in Ion Torrent data. We run SiNVICT (version 1.0) with default parameters and with no additional data pre-processing steps. We split the tumor samples (10 metastatic solid tumors plus 96 ctDNA samples) into 3 batches of similar size and we run SiNVICT simultaneously on all samples from each batch. SiNVICT applies a number of post-processing filters and calls variants at 6 different confidence levels, with level 6 assigned to variants that pass all filters. On ctDNA samples, we additionally compare AmpliSolve to deepSNV [9], a state-of-the-art method for calling low allele frequency variants in deep sequencing data (although originally designed to detect sub-clonal mutations on Illumina rather than Ion AmpliSeq data). We run deepSNV (version 1.21.3) with default parameters following the available vignette in the R package.

### How to run AmpliSolve

AmpliSolve two modules, AmpliSolveErrorEstimation and AmpliSolveVariantCalling, can be downloaded from github (https://github.com/dkleftogi/AmpliSolve). Additional requirements include running versions of the programs Samtools [15], ASEQ [16] and the Boost libraries for C++. Here we provide a brief description of how to run AmpliSolve, however, more detailed information and a number of examples are available on the github page.

For a given amplicon panel, error estimation at each genomic position, for each alternative nucleotide and for each of the two strands requires availability of amplicon-based data from N normal sample files. Although we don’t enforce a minimum value for N, values below 10 are likely to give low-quality error estimations. In general, we suggest using as many normal samples as possible when training the error matrix for your panel. If normal samples are not available, AmpliSolveErrorEstimation assigns a constant error rate to all positions, nucleotides and strands in the panel. The default constant error is 0.01 but the user can specify a different value if needed (e.g. for different sequencing platforms). Note that AmpliSolveErrorEstimation does not take bam files as input but rather bam-derived read count files. Read count files can be obtained by running the program ASEQ [16]. Once the read count files have been produced, the user needs to set the value of the C pseudo-count parameter (equation (1)). The choice of C will depend on the trade-off between precision and sensitivity the user is interested in. Users can refer to the benchmarking experiments performed in this paper. In general, values of C between 0.001 and 0.01 should suit most applications.

Once the error matrix has been calculated, it can be fed to the AmpliSolveVariantCalling program together with read count files for the tumor samples again to be produced by running ASEQ. Note that AmpliSolveVariantCalling does not require matched normal-tumor samples for calling SNVs. In fact, AmpliSolveVariantCalling calls all variants it can find in the tumor sample, including germline variants. To separate germline from somatic variants users will need to run AmpliSolveVariantCalling on a matched normal sample and take the difference between the two output files. Command-line syntax for running AmpliSolveErrorEstimation and AmpliSolveVariantCalling is provided on github.

## Results

### Sequencing error estimation, self-consistency test and AmpliSolve FDR

AmpliSolve estimates the background sequencing noise by analyzing the distribution of alternative alleles in normal samples. As previously reported, PGM errors tend to be systematic [11]. For example, we observe that A>G (T>C) and, to a lesser extent, C>T (G>A) mutations tend to have a higher background error level (see Figure S3). For this reason, AmpliSolve assigns separate error levels to each genomic position, each alternative allele and each strand (see Figure S4). These are then utilized to build the Poisson models that are at the core of AmpliSolve SNV calling (Methods). In this section, we study AmpliSolve variant calling performance as a function of two parameters: the pseudo-count *C* (equation (1) in Methods) and the number of normal samples *N* that are used to calculate the error estimations.

We perform a self-consistency test using sets of normal samples to train our models and other, non-overlapping sets of normal samples for testing them. Given a dataset of N=184 normal samples (Methods), we proceed as follows: 1) we select a number M < N of samples at random and additionally a value *c* of the C parameter; 2) we use the M samples to train our Poisson-models with C=*c*; 3) we use the models obtained in 2) to predict SNVs in the remaining N-M samples; 4) we calculate FDR and TPR by defining as negatives all alternative alleles that have VAF<20% and as positives those for which VAF≥20%. This threshold is chosen empirically based on the distributions of VAFs that we observe in the data (Figure S1); 5) we repeat steps 1) to 4) 50 times for each pair of (M,*c*) values, each time selecting a new set of M samples at random; 6) we calculate median FDR and TPR over the 50 experiments. We perform steps 1) to 6) for all combinations of the following values of M (size of the training set) and *c* (parameter C): M=10, 20, 40, 80, 120 and *c*=10^−5^, 5*10^−5^, 10^−4^, 5*10^−4^, 10^−3^, 2*10^−3^, 5*10^−3^, 10^−2^, 2*10^−2^. In Figure 2a and 2b, we plot the median FDR for each size of the training set (10-120) as a function of C; additionally, for comparison, we plot the median FDR of a method in which we skip the error estimation step and we set instead s=*c* for all positions, nucleotides and strands (‘baseline caller’) (see equation (1) in Methods for the definition of s). The FDR reported in Figure 2a is calculated by considering an SNV as called by AmpliSolve if and only if it has a Q score higher or equal 20 (i.e. p-value ≤ 0.01; this is equivalent to the SNV not having a LowQ flag, see Methods). The FDR reported in Figure 2b, instead, is calculated by considering an SNV as called by AmpliSolve if and only if the program assigns a ‘PASS’ flag to it. that is, if none of the warnings described in the Methods section applies. In Figure 2a we see that, for relatively small values of *c*, the training set size N affects the method performance, with more samples providing better error estimation and thus lower FDR. Also, our approach provides an approximately 2- to 4-fold FDR improvement over the baseline caller at all values of *c*≤0.01. For values of *c*>0.01, instead, differences with the baseline caller become negligible. Figure 2b shows that filtering AmpliSolve’s SNV calls using the warning flags that we define on top of LowQ (such as those related to low number of supporting reads, homopolymer regions, etc.) has the effect of further improving the FDR. Also, it reduces differences between FDRs obtained when using training sets of different size. All of the above findings suggest that estimating the background noise at each position, for each nucleotide and for each strand is important for reducing the number of FPs arising from noise in Ion AmpliSeq data. If we now consider the median values of the Sensitivity measure (or TPR), we discover that in all our experiments they are close to 1, irrespective of the value of M and *c*. This close to perfect Sensitivity is not surprising as our definition of positives (VAF≥20%) makes them relatively simple to discriminate from the background noise especially considering the fact that most of them have VAFs that are much higher than 20% (Figure S1). Thus, in order to truly test AmpliSolve Sensitivity, we have to perform a different kind of experiment, which we describe in the next section.

**Figure 2.**
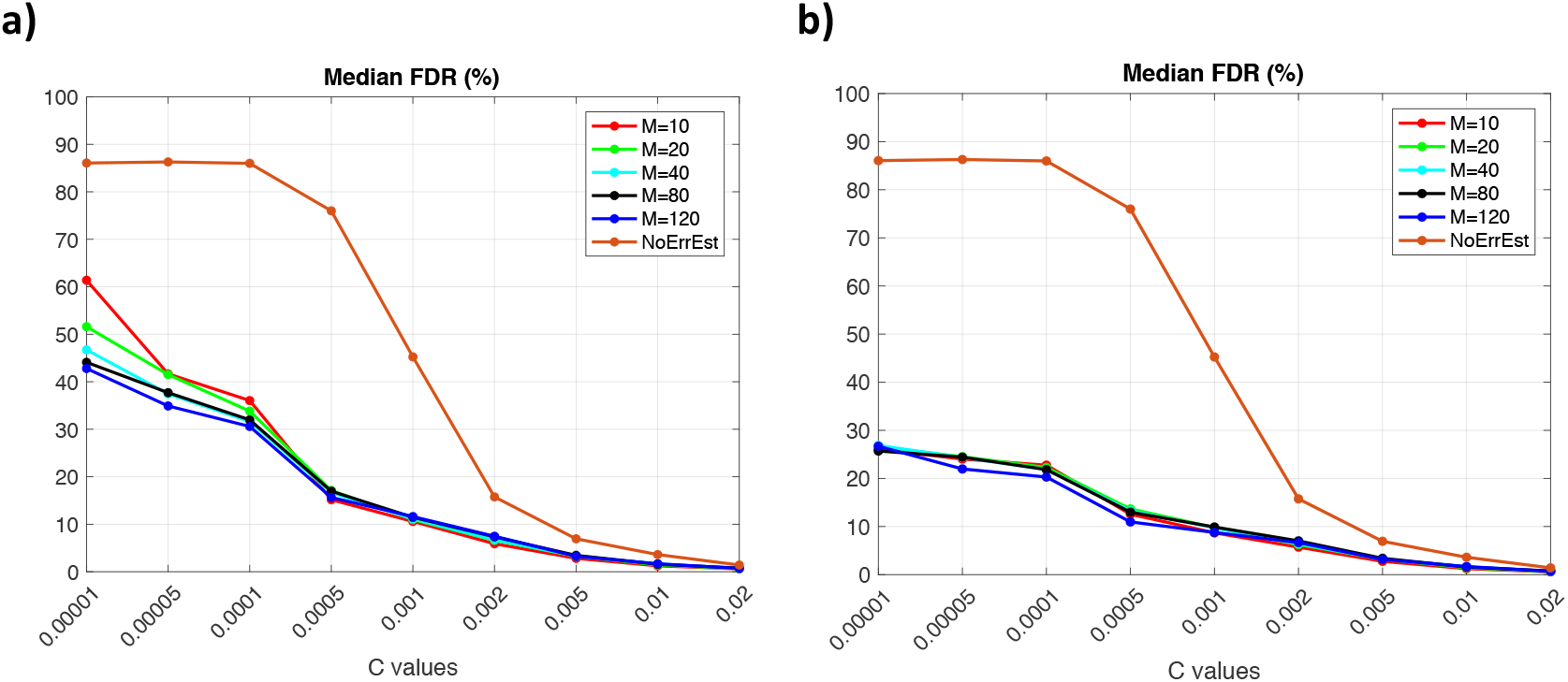
Assessing AmpliSolve’s performance using normal samples. a) Median AmpliSolve FDR (%) as a function of the model pseudo-count parameter, when using different numbers M of normal samples as training set and testing on the remaining normal samples. We consider as TP all normal variants with VAF≥20% and as FP all normal variants with VAF<20% (see Text). We consider all AmpliSolve calls that have Q-score≥20. b) Same as (a) when considering only AmpliSolve calls with a ‘PASS’ quality flag (see Text).

### Synthetic variants test for TPR estimation

In order to test the sensitivity of our method at low VAFs (0.5% to 4%), we design the following experiment. We first select two amplicons on the *AR* gene (1,017 genomic positions overall); the *AR* gene is chosen because clinically relevant but for this purpose other choices would be equally valid. Then, we use 120 normal samples randomly selected from the full set of 184 described in Methods to estimate the errors at each position in the two amplicons, for each nucleotide and each strand, according to formula (1). Next, we test the method’s sensitivity on synthetic variants. For each possible alternative allele at each of the 1,017 amplicon positions, we set read depth to a fixed value *COV* and the number of reads supporting the allele to 2*a (a* supporting reads on the forward strand and *a* on the reverse strand). We use COV=800, 1600, 3200, 6400 (values in this range apply to more than 60% of full panel positions with coverage >200, see Figure S5) and for each value of COV we select *a* corresponding to VAFs of 0.5%, 1%, 1.25%, 2%, 3% and 4%. For example for COV=800 we test *a*=2,4,5,8,12,16. We then apply the Poisson models previously trained on the 120 normal samples to predict variants at each position and for each alternative allele and consider only AmpliSolve calls with a ‘PASS’ quality flag. We consider all synthetic variants to be positives (thus, no FDR can be calculated in this case) and ask how many of these can be detected by AmpliSolve. We stress that while in each experiment the VAF is by design the same at all positions and for each alternative allele and strand, following estimation from the normal samples the error estimate is position-, alternative allele- and strand-dependent. We calculate the TPR for all combinations of *COV* and *VAF*. We do this for several values of the pseudo-count parameter C in the range of low AmpliSolve FDR as calculated from the self-consistency test in the previous section or the range of main interest for applications (C=0.001, 0.002, 0.005, 0.01, 0.02, see Figure 2b).

Figures 3(a-e) highlight the role of the C parameter as an approximate lower bound for AmpliSolve sensitivity (see equation (1)). Typically, AmpliSolve identifies few or no variants at allele frequencies equal to or lower than C, in the range of tested coverage depth (see, in particular, Figures 3c-e). For example, for C=0.005 no calls are made at VAF=0.5% even at values of COV as high a 6,400. Along the same lines, for values of C equal 0.01 and 0.02, which correspond to FDRs below 1.6% and 0.6%, respectively (Figure 2b), the lowest VAFs that AmpliSolve can detect are above 1% and 2%, respectively. For VAF values above C, on the other hand, sensitivity grows quickly with increasing VAF. For example within the depth of coverage range that we have analyzed, when using C=0.01 and C=0.02 AmpliSolve successfully calls the vast majority of synthetic variants at VAF 2% and 3%, respectively. When we compare the Sensitivity histograms in Figures 3a-e to the FDR curves in Figure 2b, we see that AmpliSolve can reliably predict synthetic SNVs at VAFs as low as 1% while still in a regime of relatively low FDR. Indeed for C=0.002, at an estimated FDR of 6.8% (Figure 2b), AmpliSolve calls most SNVs with 1% allele frequency at depth of coverage >1,600 and most SNVs with allele frequency 0.5% at depth of coverage >3,200. While it will be up to the user to select the best trade-off between FDR and TPR for a specific experiment, it would appear that values of C between 0.001 and 0.01 would likely represent a reasonable compromise between these two performance measures in most applications.

**Figure 3.**
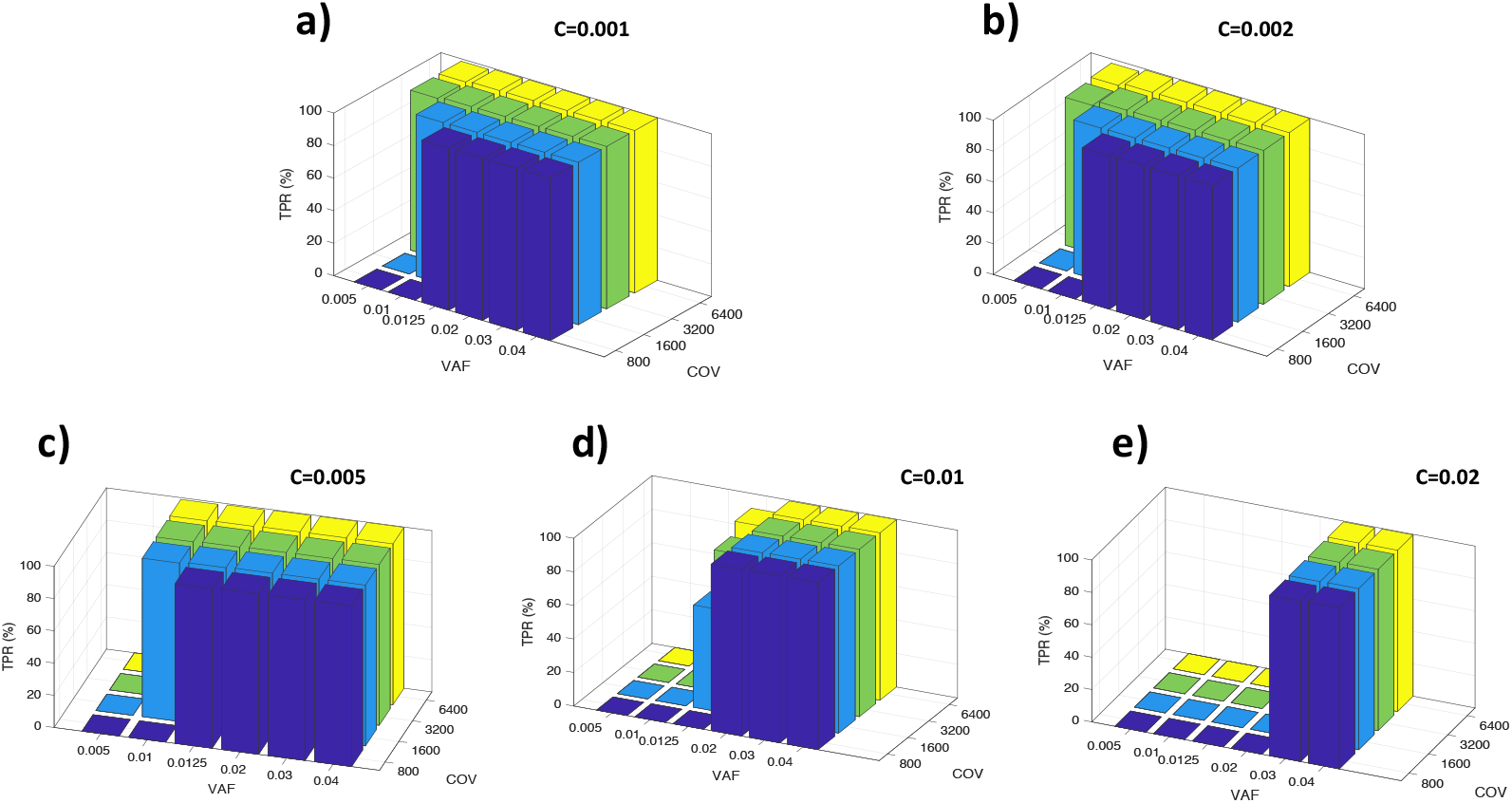
Assessing AmpliSolve’s sensitivity using synthetic data. (a-e) AmpliSolve TPR (Sensitivity) values in *in-silico* synthetic variant experiments. We test different combinations of VAF, depth of coverage and C parameter values (see Text).

### Benchmarking AmpliSolve perfomance using Illumina WGS data

For 5 additional CRPC patients, we have access to 10 metastatic solid tumor (for some patients more than one metastasis) and associated normal samples. These were sequenced both with our custom Ion AmpliSeq panel and with the Illumina platform as WGS (the latter, with average coverage ~100X) (Methods). We use these 10 samples to provide a validation of AmpliSolve SNV calls in a more realistic set-up with respect to what shown in the previous two sections. For training our AmpliSolve Poisson models, we use the full set of 184 normal samples sequenced with the Ion AmpliSeq technology and we set C=0.002.

In the solid tumor samples, when run on the Ion AmpliSeq data AmpliSolve identifies a total of 556 SNVs. For the same set of genomic positions processed by AmpliSolve, our WGS-variant calling pipeline MutPlat (Methods) calls a total of 603 SNVs in the corresponding Illumina data. The list of positions processed by AmpliSolve includes all those covered by our amplicon panel minus the ones for which no background error estimate can be produced (Methods). Almost all SNVs identified in the WGS data are germline (592 out of 603) but some of them have low VAF in the tumor samples because of deletions and loss of heterozygosity (LOH) events in the tumor DNA combined with germline DNA contamination. It is therefore a very valuable test set that includes confidently identified variants at low VAF.

The level of agreement between AmpliSolve and the ground-truth set of calls from MutPlat is summarized in Figure 4a and in Table 1. On this data, AmpliSolve achieves 87% TPR, 94% PPV and 90% F1. In particular, of the 556 SNVs called by AmpliSolve, 525 SNVs are also identified by MutPlat (TP). The remaining 31 are likely false positives (FP) although some of them might be real somatic variants with very low VAFs (and hence non detectable by a WGS done at 100x). The 78 SNVs additionally identified by MutPlat in the WGS data are likely AmpliSolve false negatives (FP). However, we note that 48 of them correspond to positions not called because the coverage in the tumor samples was below the threshold of 100 reads per strand and 26 were filtered out because of the strandBias flag. By simply setting more lenient parameters RD_min_=50 and SB_th_=0.01 we are able to drastically reduce the number of missed calls without affecting AmpliSolve precision. With these settings we obtain 571 TP, 34 FP and 32 FN, which translates into 95% TPR, 94% PPV and 94% F1.

**Figure 4.**
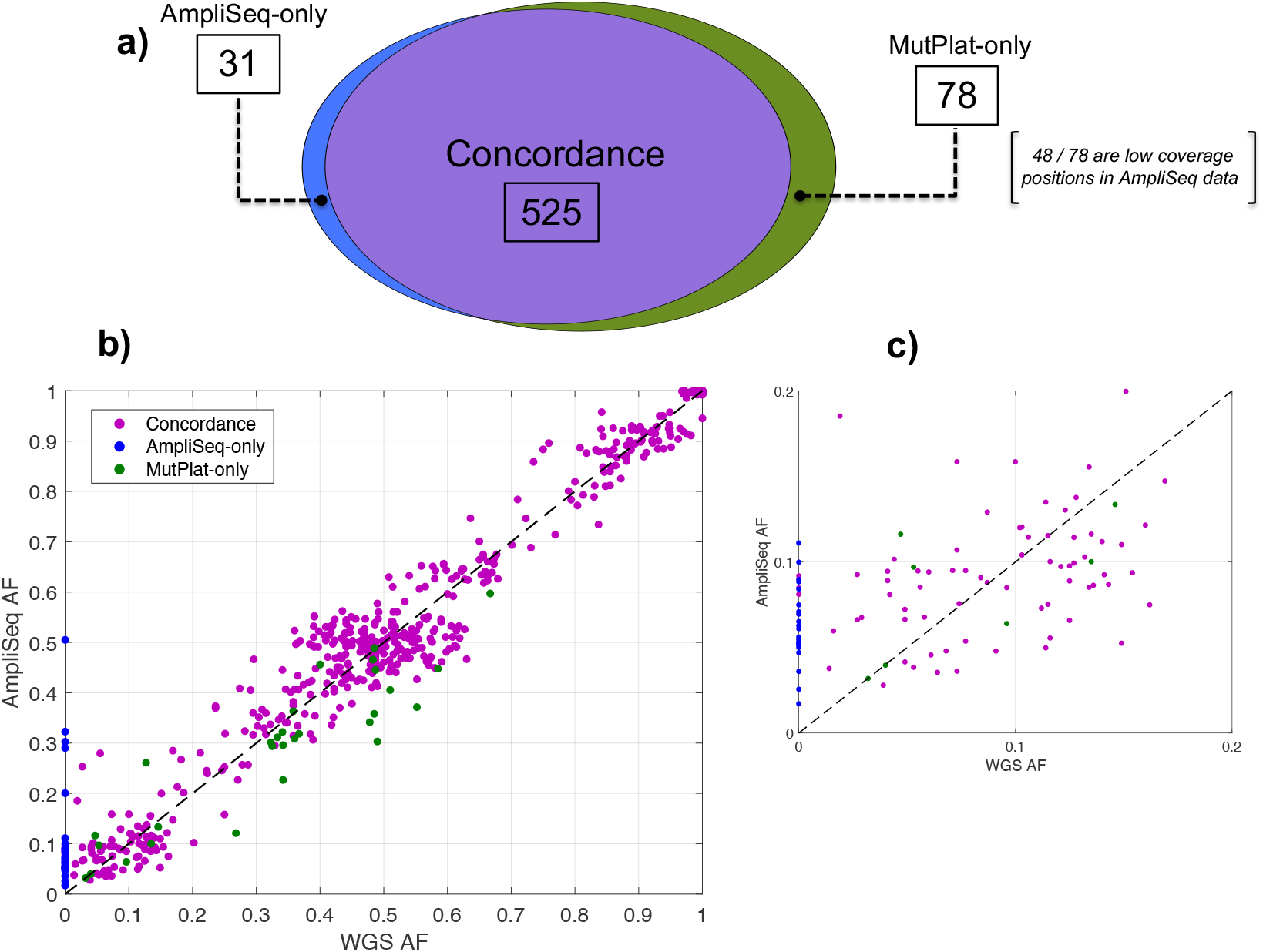
Benchmarking AmpliSolve calls with Illumina WGS calls. a) Venn diagram of mutations on 10 samples sequenced with both Ion Torrent and Illumina platforms and called respectively by AmpliSolve and by MutPlat. Low coverage positions denote mutations excluded by AmpliSolve because poorly covered (<100 reads on at least one strand, ‘uncallable’ by AmpliSolve). (b) Scatter plot of VAFs in WGS and AmpliSeq data. Note that all the SNVs not called by AmpliSolve (green point) have some support in the data and are reported in its output (hence they have AF > 0) but are filtered out, mostly because of strand bias. (c) Same as (b) but for VAFs<20%. Note that some concordant calls (purple points) have WGS AF=0; these are real germline variants with no support in the tumor (Methods). For the sake of this comparison, both in (b) and in (c) we don’t consider the 49 mutations at positions of low coverage in Ion Ampliseq data (see (a)) (‘uncallable’ for AmpliSolve).

**Table 1.**
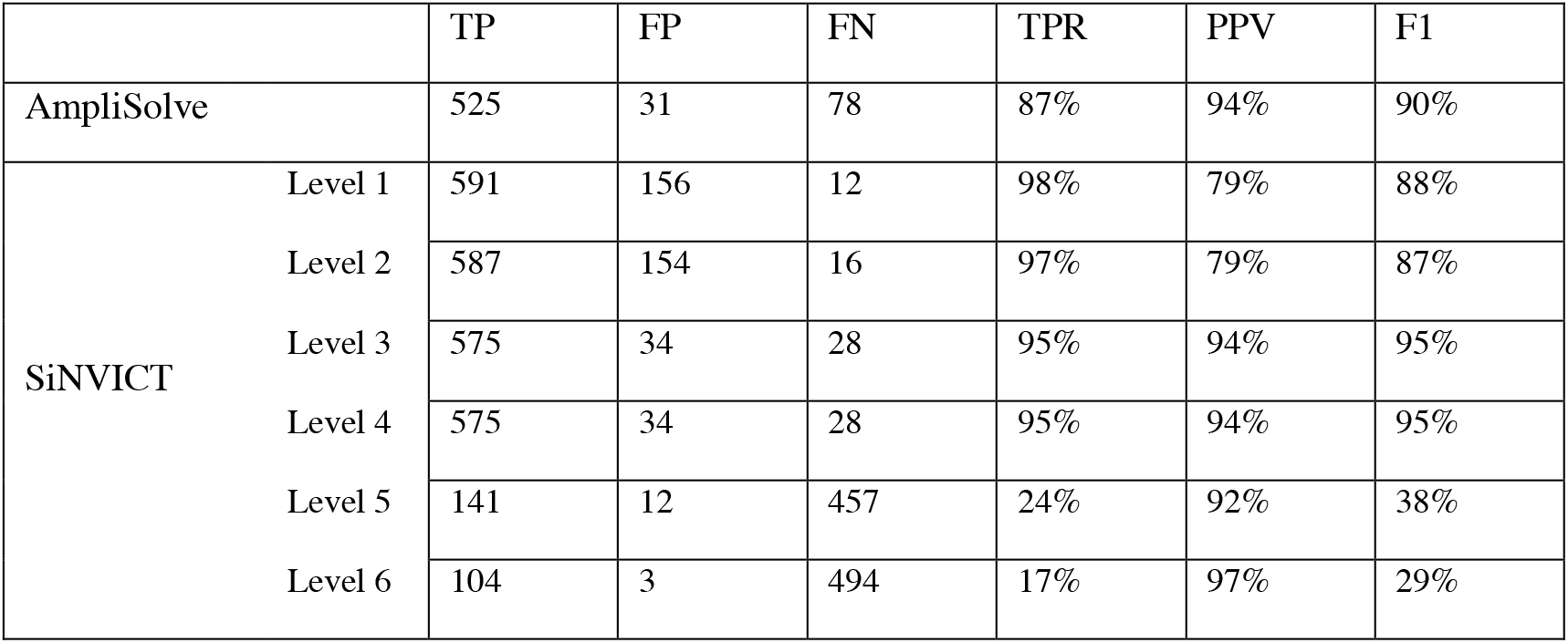
Comparison between AmpliSolve and SiNVICT calls across the targeted panel. MutPlat calls on Illumina WGS data have been used as ground-truth. SiNVICT levels correspond to confidence levels in the calls (6 being the highest). TP=True Positives, FP=False Positives, FN= False Negatives, TPR=True Positives Rate (Sensitivity), PPV=Positive Predictive Value (Precision), F1=Harmonic mean of Precision and Sensitivity.

In Figure 4b and 4c we report a scatter plot of the VAFs in the WGS and AmpliSeq data with colors indicating common calls (purple), MutPlat-only calls (green) and AmpliSolve-only calls (blue), respectively. Overall there is a good concordance between AmpliSolve and MutPlat calls, even at low VAF (Figure 4c). In particular, AmpliSolve correctly identifies 18 out of 21 SNVs with VAF < 5% in the WGS calls. AmpliSolve does call a number of likely false positives at low VAF, however we note that most of them occur at recurrent positions across patients and could therefore potentially be identified and discarded at a post variant calling analysis stage.

In Table 1, we additionally compare AmpliSolve’s performance to the one of SiNVICT when run on the same 10 solid tumor samples.(MutPlat calls on the Illumina data are used as ground truth in both cases). SiNVICT assigns a confidence level (1 to 6) to its calls according to a series of hierarchical filters (each filter eliminates some calls from the previous level). On our dataset, SINVICT’s highest confidence level (level 6) although very precise appears to miss a substantial number of SNVs (i.e. it has low sensitivity), especially at low VAF. Better overall results are obtained at confidence levels 3 and 4, In this case, precision and sensitivity values are similar to the one obtained by Amplisolve when using RD_min_=50 and SB_th_=0.01. Interestingly, at low VAF AmpliSolve and SiNVICT seem to identify slightly different sets of SNVs, suggesting that it might be possible to improve SNV calling by appropriately combining them.

### Clinical application using ctDNA samples and ddPCR for validation

One of the most promising clinical applications of ctDNA is profiling of specific mutations associated with tumor progression and resistance to cancer therapies. To evaluate AmpliSolve’s usefulness for this important task, we use results from a ddPCR screen on 96 samples from our CRPC patients at three genomic positions within the *AR* gene, which are associated with resistance to targeted therapy (Methods). ddPCR detects 30 variants in total at these positions in a VAF range of 0.1 to 49% (note, however, that in some experiments only the presence or absence of the variant was recorded). Next, we compare AmpliSolve calls at the same positions in the AmpliSeq NGS data for the same samples (predictions made after training AmpliSolve with pseudo-count parameter C=0.002 on 184 normal samples). In Figure 5a we summarize the level of agreement between AmpliSolve and the ddPCR experiments. AmpliSolve correctly calls 19 out of 30 ddPCR variants and predicts variants at two additional positions. If we take the ddPCR experiments as our ground-truth, this translates into 90% PPV 63% TPR and 74% F1 for AmpliSolve at these 3 clinically relevant genomic positions.

**Figure 5.**
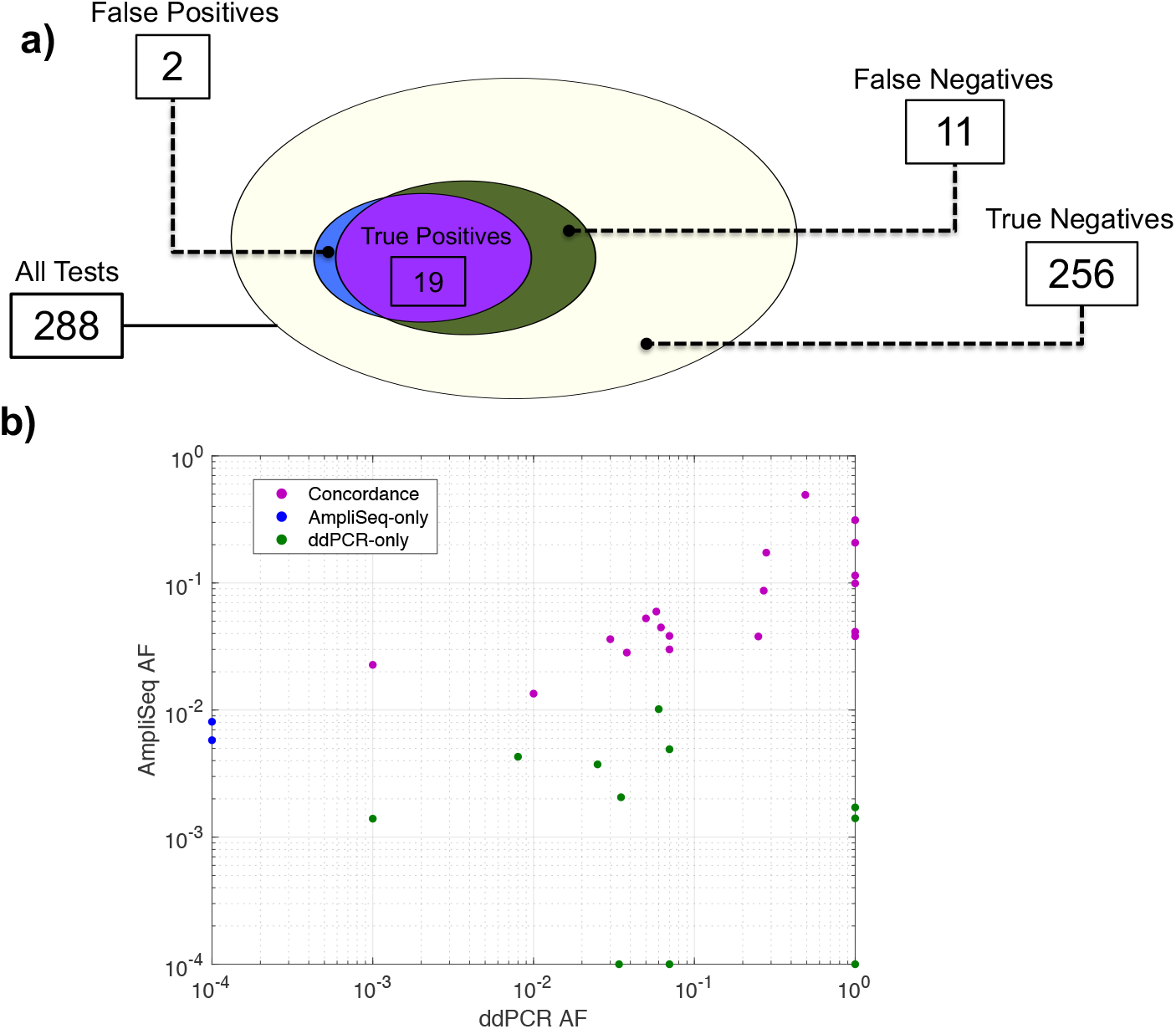
Validating AmpliSolve performance with ddPCR experiments. a) Venn diagram of mutations in 96 samples at 3 positions as determined by AmpliSolve and ddPCR experiments. False positives refer to variants called by AmpliSolve and not detected by ddPCR, false negatives the opposite. In 256 out 288 cases neither AmpliSolve nor ddPCR detect a mutation. (b) Scatter plot of the VAFs in the ddPCR and Ion Torrent data. Most of the SNVs missed by AmpliSolve (green points) have some support in the NGS data but they cannot be distinguished from noise. Because of the log scale, we arbitrarily set AF=10-4 for negative calls with AF=0. Similarly, we set AF=1 for ddPCR calls for which no allele frequency information is available.

As a comparison, we run the SiNVICT and deepSNV methods on the same Ion Torrent data and extract their SNV calls at these positions. The results are summarized in Table 2. Similar to what observed in the previous section, SiNVICT highest confidence levels (5 and 6) have low sensitivity (30% TPR). Better results are obtained at lower confidence level (1 to 4) whereby SiNVICT correctly identifies 17 out of 30 ddPCR variants without introducing any false positives. In total deepSNV calls 18 SNVs, 15 of which are correct.

**Table 2.**
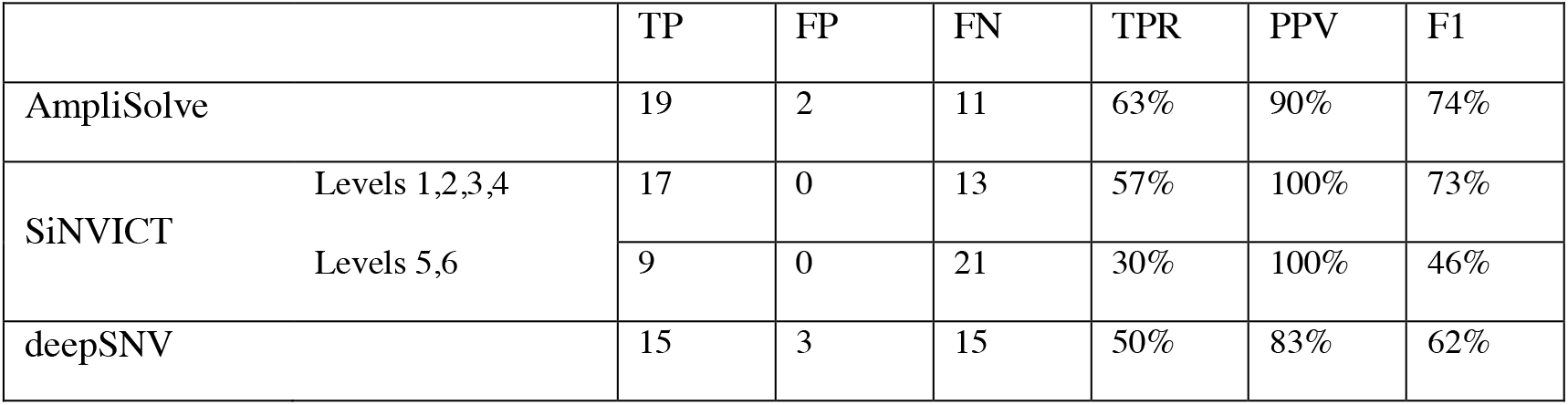
Comparison of SNV calling on 96 samples at 3 genomics positions. The 3 positions on the AR gene were screened by ddPCR used here as ground-truth. SINVICT Levels 5 and 6 and Levels 1, 2, 3 and 4 have been grouped as they give the same results. TP=True Positives, FP=False Positives, FN= False Negatives, TPR=True Positives Rate (Sensitivity), PPV=Positive Predictive Value (Precision), F1=Harmonic mean of Precision and Sensitivity.

When looking at AmpliSolve predictions in more details (Figure 5b), we note that all ddPCR positives not called by our program (and additionally missed by both SiNVICT and deepSNV) have VAF<1% in the NGS data and that AmpliSolve succeeds in calling all ddPCR positives at higher NGS frequencies including several at VAFs between 1% and 5%. It is also important to note that AmpliSolve correctly predicts 256 out of 258 (99.2%) ddPCR negatives. While the results presented in this section refer to only three genomic positions, they are indicative of AmpliSolve’s potential value in a clinically relevant setting.

## Discussion

In this study we present AmpliSolve, a new bioinformatics method that combines position-specific, nucleotide-specific and strand-specific background error estimation with statistical modeling for SNV detection in amplicon-based deep sequencing data. AmpliSolve is originally designed for the Ion AmpliSeq platform that is affected by higher error levels compared to, for example, Illumina platforms. Our method is based on the estimation of noise levels from normal samples and uses a Poisson model to calculate the p-value of the detected variant. We assess AmpliSolve’s performance with experiments that use normal samples (self-consistency tests) and simulated data (synthetic variants) and, additionally, with tests that utilize real metastatic samples sequenced with both Ion Torrent and Illumina platforms. In these experiments, AmpliSolve achieves a good balance between precision and sensitivity, even at VAF < 5%. These experiments also suggest possible ways to further improve the method, such as adopting a better strand bias filter, reducing the minimum coverage requirement for calling a variant and introducing a ‘black list’ of positions characterized by an unusual noise distribution across samples (e.g. bimodal). Further, we test AmpliSolve in a clinical relevant setting by calling SNVs in 96 liquid biopsy samples at 3 positions that had been additionally screened by ddPCR assay. In this experiment AmpliSolve successfully identifies SNVs at VAF as low as 1% in the NGS data. This opens up interesting possibilities for clinical applications using the Ion Torrent PGM such as, for example, tracking mutations in ctDNA to monitor treatment effectiveness and/or disease relapse.

## Conclusions

AmpliSolve is a new computational tool for the detection of low frequency SNVs in targeted deep sequencing data. It uses a set of germline samples to build a sequencing error profile at each genomic position of interest. Based on these profiles AmpliSolve estimates the likelihood of a variant being real or just the result of sequencing artefacts. We test AmpliSolve on clinical cancer samples sequenced with a custom Ion AmpliSeq gene panel and show that AmpliSolve can correctly identify variants even at allele frequency below 5% and as low as 1%. This is significant because detecting variants with low allele frequency can be challenging using Ion Torrent sequencing. From a methodological point of view, we believe that the use of models with position-specific error estimates, as described here, could have a significant impact on variant detection for other sequencing platforms as well.

## List of abbreviations

NGS: Next-Generation Sequencing
SNV: Single Nucleotide Variant
VAF: Variant Allele Frequency
cfDNA: circulating free DNA
ctDNA: circulating tumor DNA
PGM: Personal Genome Machine
CRPC: Castration Resistant Prostate Cancer
WGS: Whole Genome Sequencing
ddPCR: digital droplet PCR
TP: True Positives
FP: False Positives
FN: False Negatives
TPR: True Positive Rate
PPV: Positive Predictive Value
FDR: False Discovery Rate

## Declarations

### Ethics approval and consent to participate

All samples were collected in studies with institutional regulatory board approval and conducted in accordance with the Declaration of Helsinki and the Good Clinical Practice guidelines of the International Conference of Harmonization (REC numbers: 04/Q0801/6 at the Royal Marsden, London, UK, 2192/2013 at the Istituto Scientifico Romagnolo per lo Studio e la Cura dei Tumori, Meldola, Italy and 15/98 at Peter MacCallum Cancer Centre). Written informed consent was obtained from all patients.

### Consent for publication

Patients have consented that next-generation sequencing data with no personal identifiers can be used for publication in an anonymized format.

### Availability of data and material

AmpliSolve is freely available at https://github.com/dkleftogi/AmpliSolve. The datasets analysed during the current study is available from the authors on reasonable request.

### Authors’ contributions

DK implemented and tested the AmpliSolve software. DK, MP and SL designed the computational study and analyzed the results. SS, SQW, DW, AJ, VC and DGT were responsible for sample collection; DW, AJ, VC and DGT carried out library preparation and DNA sequencing. DW, AJ, VC and DGT performed the ddPCR assays. SL and GA conceived the study; SL, GA and MP supervised it. DK, MP and SL wrote the paper, DW and GA reviewed it. All authors read and approved the final manuscript.

## Acknowledgements

The authors thank the participating men and their families who suffered from metastatic prostate cancer and nonetheless gave the gift of participation so that others might benefit.

## Additional Files

Additional file 1: Supplementary Methods. Details of the MutPlat pipeline.

Given a set of bam files, corresponding to N tumour samples and a germline from the same patient (bamT◻◻◻ bamT◻, …, bamT◻◻◻ bamGL), the MutPlat pipeline implemented in this work consists of the following basic steps:

1. Run Mutect2 (GATK version 3.6) with default parameters on each tumour/germline pair (bamT◻◻◻bamGL◻◻ bamT◻/bamGL, …, bamT◻◻bamGL). This generates N vcf files (vcfT◻◻◻ vcfT◻, …, vcfT◻◻.
2. From the N vcf files above, generate a single vcf file, vcfT◻◻which overlays all positions present in the individual vcf files. As an example if vcfT◻◻= {pos1, pos2, pos3} and vcfT◻◻= {pos2, pos4} then vcfT◻= {pos1, pos2, pos3, pos4}. All positions in the individual vcf files are included, independently of e.g. coverage and filter flag assigned by Mutect2.
3. Run Platypus (version 0.8.1) jointly on (bamT◻◻◻ bamT◻, …, bamT◻◻◻ bamGL) using the vcf file, vcfT, from step 2 as prior. More specifically we set the options ‘--source=vcfT’ and ‘--getVariantsFromBam=1’.

The outcome is a multi-sample vcf file that is subsequently filtered as described in the Methods section to extract germline and somatic SNVs.

**Additional file 2: Figure S1.**
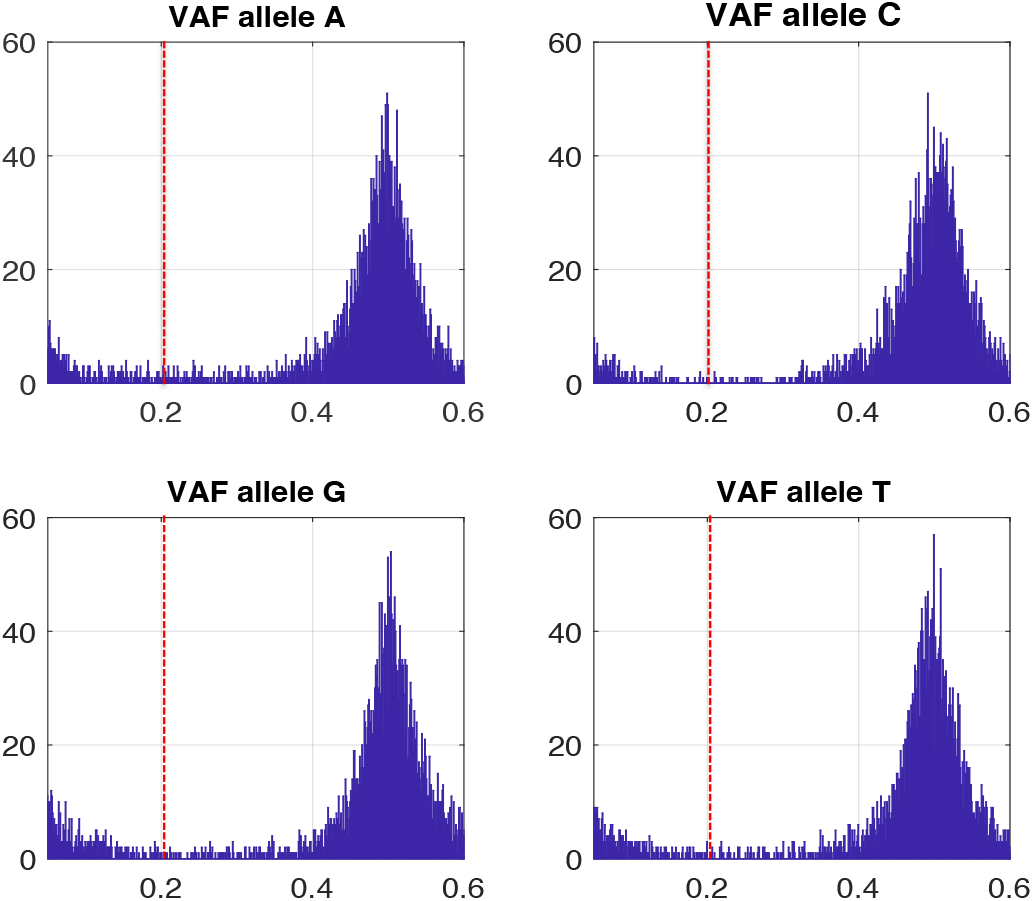
Variant allele frequency (VAF) distributions for the A, T, C, G nucleotides as calculated from 30 randomly chosen normal samples across our custom AmpliSeq panel. Only VAFs < 60% are displayed. The red lines marks VAF = 20%.

**Additional file 3: Figure S2.**
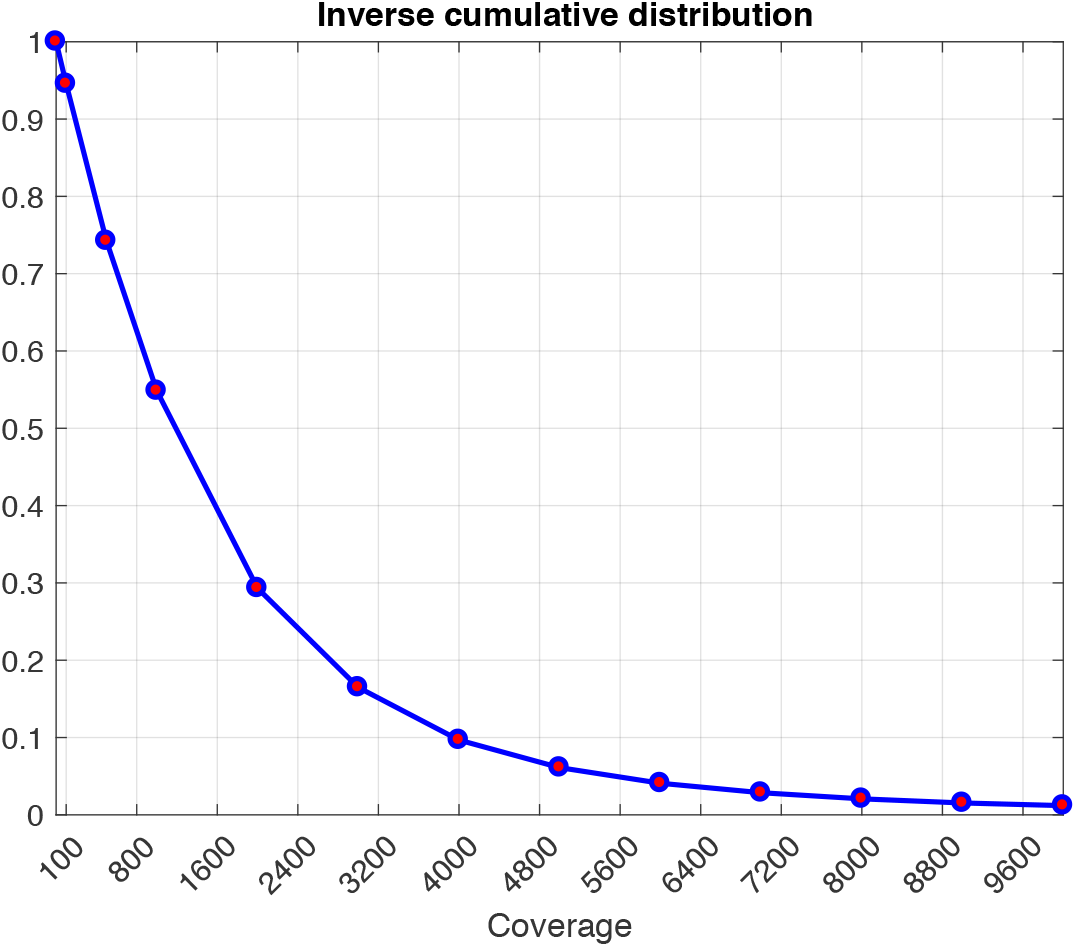
Fraction of sites in a normal sample sequenced at a given coverage or more across our custom AmpliSeq panel. The values are calculated over 30 random samples

**Additional file 4: Figure S3.**
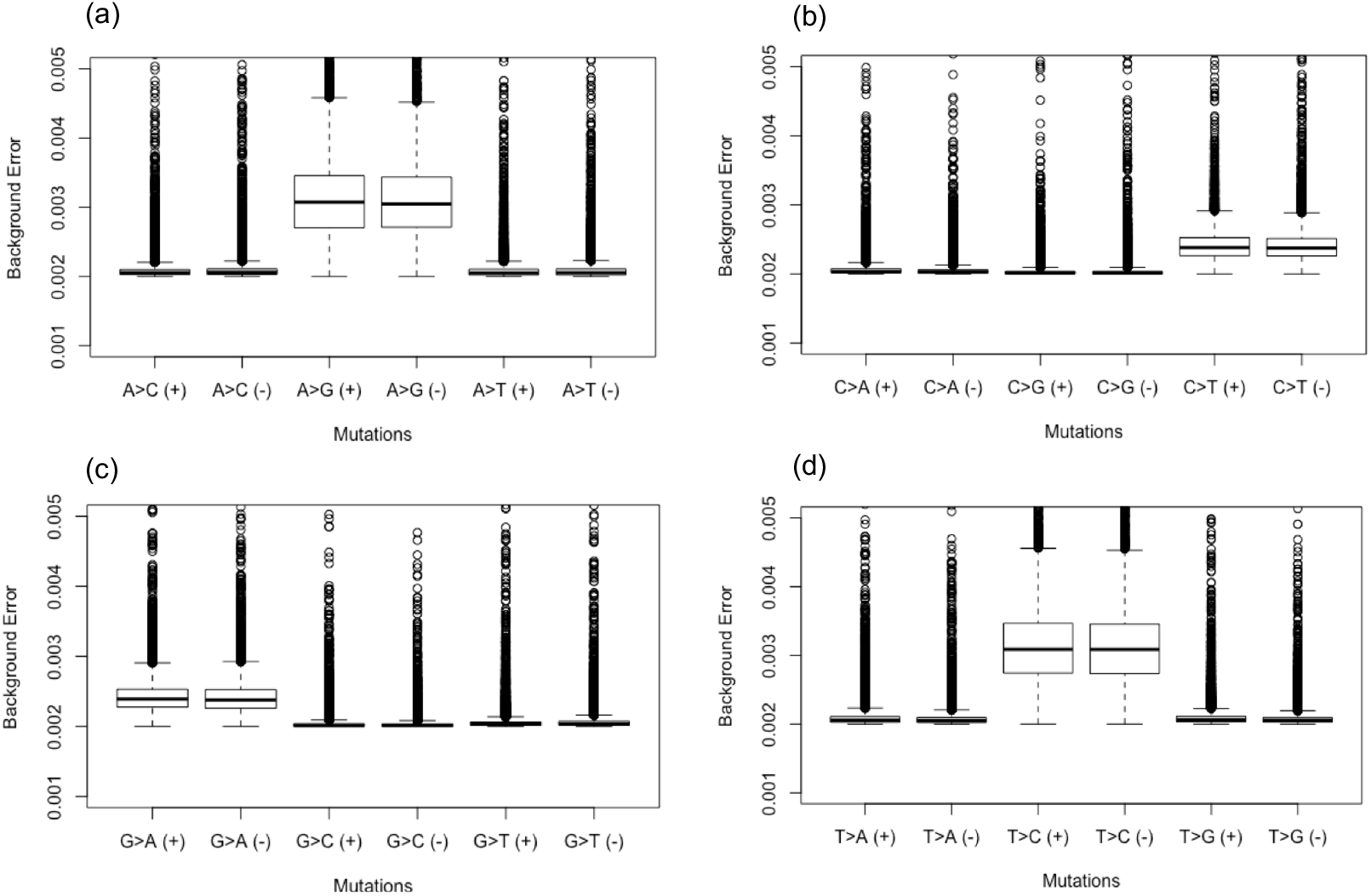
Distributions of background error values by mutation type. Panels (a), (b), (c) and (d) refers to mutations from reference allele A, C, G and T respectively. Mutations are split by alternative allele and strand, (+) and (−). Note the higher error values for A>G (T>C) and C>T (G>A) mutations. Plots are bound to error values of 0.005 on the y-axis for visual clarity.

**Additional file 5: Figure S4.**
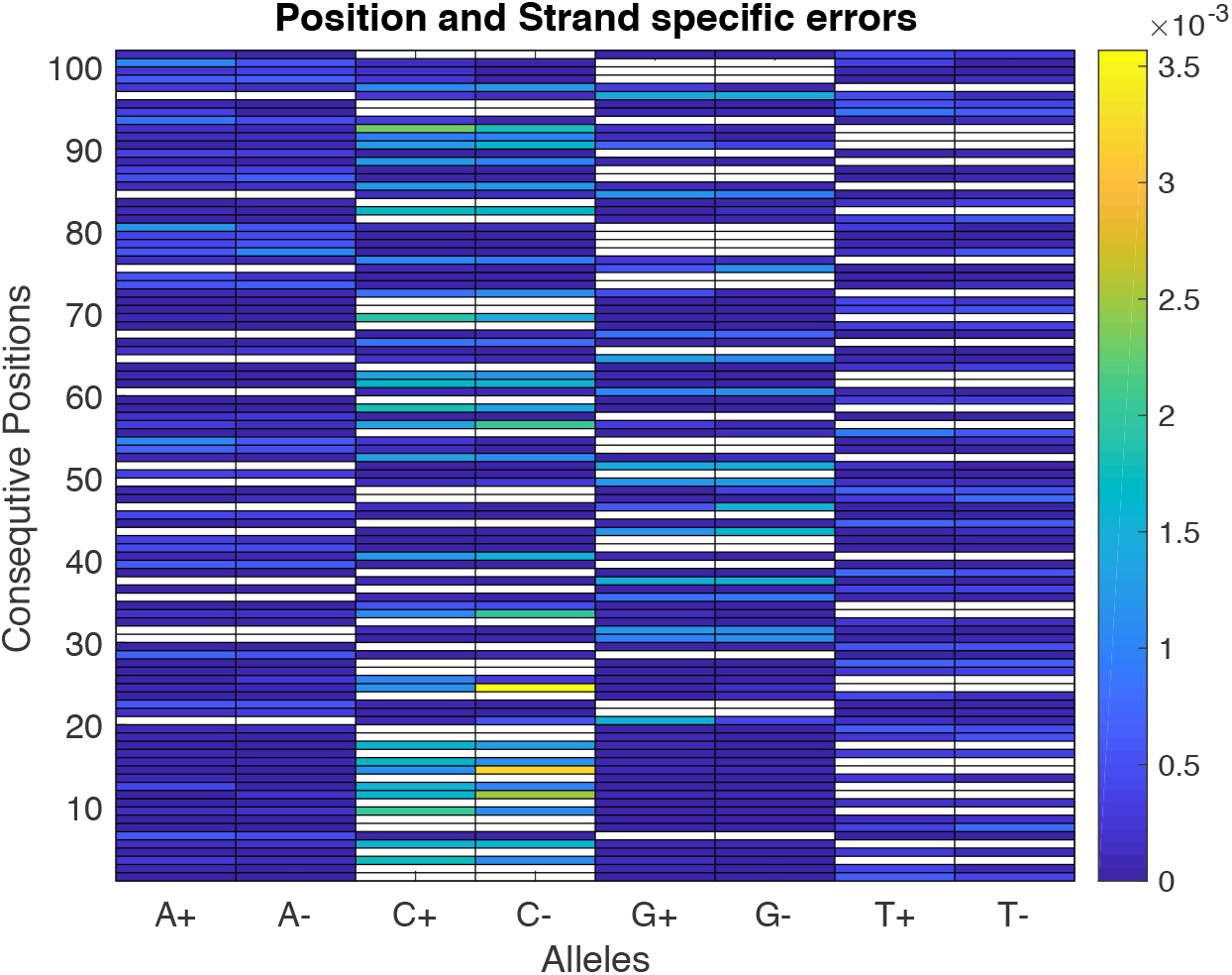
Position-specific, allele-specific and strand-specific frequency of alternative alleles in 100 consecutive positions in the *AR* gene.

**Additional file 6: Figure S5.**
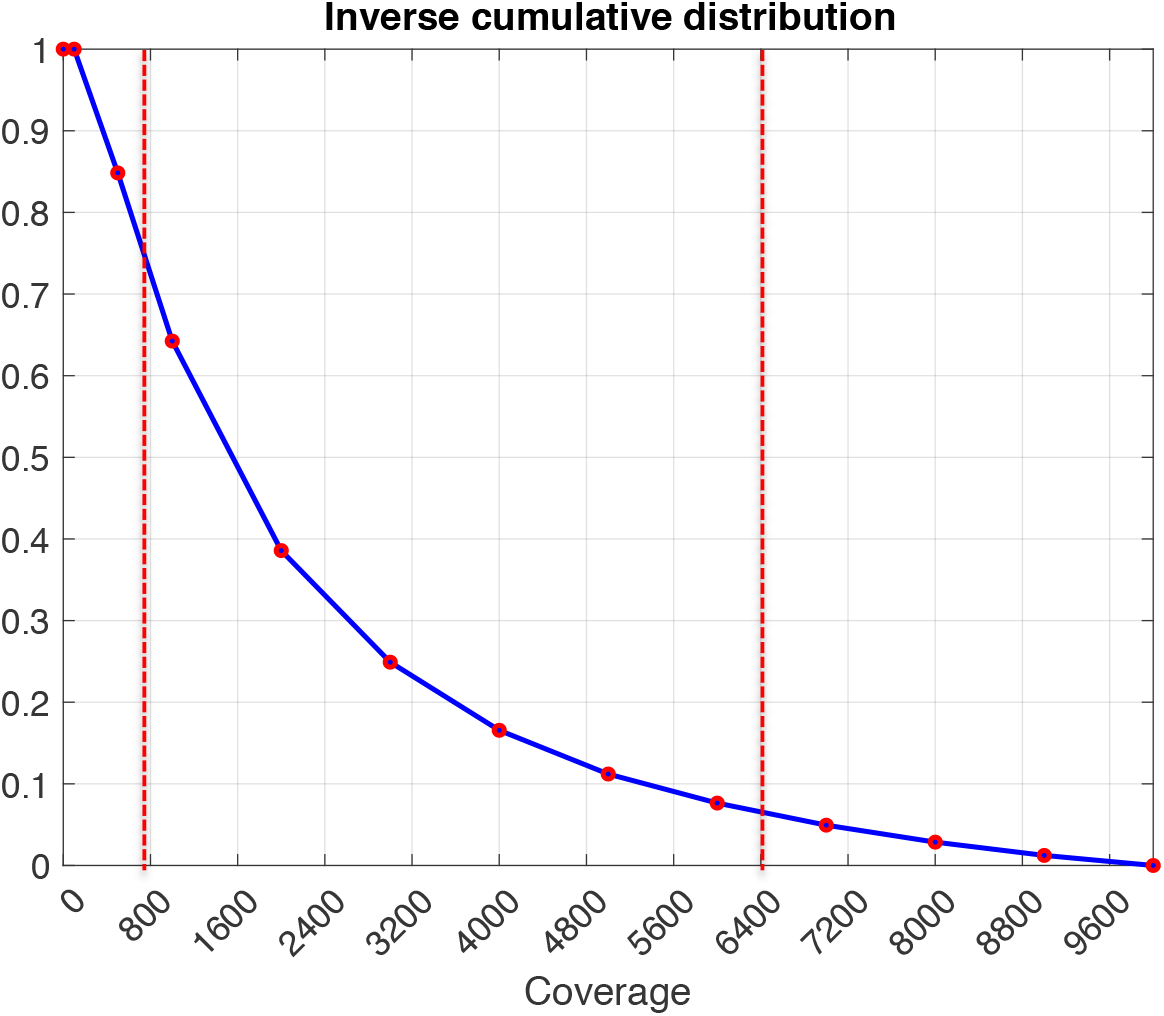
Fraction of sites in our custom AmpliSeq panel sequenced at a given coverage or more. The values are calculated over 30 randomly selected ctDNA samples. Note that positions with depth of coverage less than 200 are not considered for calculating the total number of positions. The red lines represent the upper and lower bounds of coverage used in the synthetic variant test.

## References

1. Goodwin S, McPherson JD, McCombie WR. Coming of age: ten years of next-generation sequencing technologies. Nat Rev Genet. 2016;17:333–51.

2. McGranahan N, Swanton C. Clonal Heterogeneity and Tumor Evolution: Past, Present, and the Future. Cell. 2017;168:613–28.

3. Heitzer E, Perakis S, Geigl JB, Speicher MR. The potential of liquid biopsies for the early detection of cancer. NPJ Precis Oncol. 2017;1:36.

4. Wan JCM, Massie C, Garcia-Corbacho J, Mouliere F, Brenton JD, Caldas C, et al. Liquid biopsies come of age: towards implementation of circulating tumour DNA. Nat Rev Cancer. 2017;17:223–38.

5. Newman AM, Lovejoy AF, Klass DM, Kurtz DM, Chabon JJ, Scherer F, et al. Integrated digital error suppression for improved detection of circulating tumor DNA. Nat Biotechnol. 2016;34:547–55.

6. Mansukhani S, Barber LJ, Kleftogiannis D, Moorcraft SY, Davidson M, Woolston A, et al. Ultra-Sensitive Mutation Detection and Genome-Wide DNA Copy Number Reconstruction by Error-Corrected Circulating Tumor DNA Sequencing. Clin Chem. 2018;64:1626–35.

7. Nielsen R, Paul JS, Albrechtsen A, Song YS. Genotype and SNP calling from next-generation sequencing data. Nat Rev Genet. 2011;12:443–51.

8. Xu C. A review of somatic single nucleotide variant calling algorithms for next-generation sequencing data. Comput Struct Biotechnol J. 2018;16:15–24.

9. Gerstung M, Beisel C, Rechsteiner M, Wild P, Schraml P, Moch H, et al. Reliable detection of subclonal single-nucleotide variants in tumour cell populations. Nat Commun. 2012;3:811.

10. Quail MA, Smith M, Coupland P, Otto TD, Harris SR, Connor TR, et al. A tale of three next generation sequencing platforms: comparison of Ion Torrent, Pacific Biosciences and Illumina MiSeq sequencers. BMC Genomics. 2012;13:341.

11. Bragg LM, Stone G, Butler MK, Hugenholtz P, Tyson GW. Shining a light on dark sequencing: characterising errors in Ion Torrent PGM data. PLoS Comput Biol. 2013;9:e1003031.

12. Kockan C, Hach F, Sarrafi I, Bell RH, McConeghy B, Beja K, et al. SiNVICT: ultra-sensitive detection of single nucleotide variants and indels in circulating tumour DNA. Bioinformatics. 2017;33:26–34.

13. Deshpande A, Lang W, McDowell T, Sivakumar S, Zhang J, Wang J, et al. Strategies for identification of somatic variants using the Ion Torrent deep targeted sequencing platform. BMC Bioinformatics. 2018;19:5.

14. Shin S, Lee H, Son H, Paik S, Kim S. AIRVF: a filtering toolbox for precise variant calling in Ion Torrent sequencing. Bioinformatics. 2018;34:1232–4.

15. Li H, Handsaker B, Wysoker A, Fennell T, Ruan J, Homer N, et al. The Sequence Alignment/Map format and SAMtools. Bioinformatics. 2009;25:2078–9.

16. Romanel A, Lago S, Prandi D, Sboner A, Demichelis F. ASEQ: fast allele-specific studies from next-generation sequencing data. BMC Med Genomics. 2015;8:9.

17. Romanel A, Gasi Tandefelt D, Conteduca V, Jayaram A, Casiraghi N, Wetterskog D, et al. Plasma AR and abiraterone-resistant prostate cancer. Sci Transl Med. 2015;7:312re10.

18. Rimmer A, Phan H, Mathieson I, Iqbal Z, Twigg SRF, WGS500 Consortium, et al. Integrating mapping-, assembly- and haplotype-based approaches for calling variants in clinical sequencing applications. Nat Genet. 2014;46:912–8.

19. Carreira S, Romanel A, Goodall J, Grist E, Ferraldeschi R, Miranda S, et al. Tumor clone dynamics in lethal prostate cancer. Sci Transl Med. 2014;6:254ra125.

20. Lawrence MG, Obinata D, Sandhu S, Selth LA, Wong SQ, Porter LH, et al. Patient-derived Models of Abiraterone- and Enzalutamide-resistant Prostate Cancer Reveal Sensitivity to Ribosome-directed Therapy. Eur Urol. 2018;74:562–72.

21. Conteduca V, Wetterskog D, Sharabiani MTA, Grande E, Fernandez-Perez MP, Jayaram A, et al. Androgen receptor gene status in plasma DNA associates with worse outcome on enzalutamide or abiraterone for castration-resistant prostate cancer: a multi-institution correlative biomarker study. Ann Oncol. 2017;28:1508–16.

22. Barry P, Vatsiou A, Spiteri I, Nichol D, Cresswell GD, Acar A, et al. The spatio-temporal evolution of lymph node spread in early breast cancer. Clin Cancer Res. 2018;:clincanres.3374.2018.

23. Jiang H, Lei R, Ding S-W, Zhu S. Skewer: a fast and accurate adapter trimmer for next-generation sequencing paired-end reads. BMC Bioinformatics. 2014;15:182.

24. Li H. Aligning sequence reads, clone sequences and assembly contigs with BWA-MEM. arXiv:13033997 [q-bio]. 2013. http://arxiv.org/abs/1303.3997. Accessed 29 Oct 2018.

25. Picard: A set of command line tools (in Java) for manipulating high-throughput sequencing (HTS) data and formats such as SAM/BAM/CRAM and VCF. http://broadinstitute.github.io/picard/. Accessed 30 October 2018.

26. Cibulskis K, Lawrence MS, Carter SL, Sivachenko A, Jaffe D, Sougnez C, et al. Sensitive detection of somatic point mutations in impure and heterogeneous cancer samples. Nat Biotechnol. 2013;31:213–9.

27. IGSR: The International Genome Sample Resource. http://www.internationalgenome.org/. Accessed 30 October 2018

